# acI Actinobacteria Assemble a Functional Actinorhodopsin with Natively-synthesized Retinal

**DOI:** 10.1101/367102

**Authors:** Jeffrey R. Dwulit-Smith, Joshua J. Hamilton, David M. Stevenson, Shaomei He, Ben O. Oyserman, Francisco Moya-Flores, Daniel Amador-Noguez, Katherine D. McMahon, Katrina T. Forest

## Abstract

Freshwater lakes harbor complex microbial communities, but these ecosystems are often dominated by acI Actinobacteria from three clades (acI-A, acI-B, acI-C). Members of this cosmopolitan lineage are proposed to bolster heterotrophic growth using phototrophy because their genomes encode actino-opsins (*actR*). This model has been difficult to experimentally validate because acI are not consistently culturable. In this study, using genomes from single cells and metagenomes, we provide a detailed biosynthetic route for many acI-A and -B members to synthesize retinal and its carotenoid precursors. Accordingly, these acI should be able to natively assemble light-driven actinorhodopsins (holo-ActR) to pump protons, in contrast to acI-C members and other bacteria that encode opsins but lack retinal-production machinery. Moreover, we show that all acI clades contain genes for a complex carotenoid pathway that starts with retinal precursors. Transcription analysis of acI in a eutrophic lake shows that all retinal and carotenoid pathway operons are transcribed and that *actR* is among the most highly-transcribed of all acI genes. Furthermore, heterologous expression of retinal pathway genes shows that lycopene, retinal, and ActR can be made. Model cells producing ActR and the key acI retinal-producing β-carotene oxygenase formed acI-holo-ActR and acidified solution during illumination. Our results prove that acI containing both ActR and retinal-production enzymes have the capacity to natively synthesize a green light-dependent outward proton-pumping rhodopsin.

**IMPORTANCE:** Microbes play critical roles in determining the quality of freshwater ecosystems that are vital to human civilization. Because acI Actinobacteria are ubiquitous and abundant in freshwater lakes, clarifying their ecophysiology is a major step in determining the contributions that they make to nitrogen and carbon cycling. Without accurate knowledge of these cycles, freshwater systems cannot be incorporated into climate change models, ecosystem imbalances cannot be predicted, and policy for service disruption cannot be planned. Our work fills major gaps in microbial light utilization, secondary metabolite production, and energy cycling in freshwater habitats.

## INTRODUCTION

Fresh water lakes contain complex and dynamic microbial communities that influence water quality by mediating biogeochemical cycling. These ecosystems are often numerically dominated by three clades of acI Actinobacteria (acI-A, acI-B, acI-C), ultra-microbacteria with low GC genomes that are streamlined to approximately 1.2-1.5 MBp (1-5). The ubiquity and abundance of acI raise the question, what are the factors that enable their success? Major proposed advantages include scavenging resources with membrane transporters (3, 6), evading protist feeding by virtue of their small size (7, 8), and supplementing energy needs via photoheterotrophy. Light utilization is feasible because each clade encodes actino-opsin (ActR), the putative platform protein for actinorhodopsin (holo-ActR) (3, 9–11). Validating this hypothesis not only requires a demonstration that acI can synthesize or acquire sufficient retinal, the chromophore needed to complete holo-ActR, but also biochemical evidence that the holo-ActR is functional. Additionally, the relevant genes must be shown to be expressed in the native freshwater environments where acI is dominant. That these requirements are satisfied by acI is not guaranteed; *Rhodoluna lacicola* holo-ActR was shown to be active if, and only if, exogenous retinal was added (12). Without an encoded, retinal-producing enzyme or a solution for obtaining dilute environmental chromophores, it is unclear what the physiological role and ecosystem implications of ActR in such organisms are. On the other hand, the Luna1 bacterium *Candidatus* Rhodoluna planktonica encodes a putative retinal-synthesizing enzyme (13). This family of Actinobacteria is radically different from acI and much less abundant and prevalent in freshwater lakes (2), but this organism provides motivation to determine if acI truly use rhodopsins for photoheterotrophy.

Rhodopsins are seven-transmembrane-helix bundles that bind retinal at an internal lysine via a Schiff base (14). The retinal pocket amino acid sidechains, specifically a tuning residue (15), determine the maximal absorption wavelength of the holo-protein. The captured photon energy triggers the geometric isomerization of retinal, which completes a transfer pathway through the interior of the bundle and thus across the membrane. Most commonly, protons are shuttled out of the cytoplasm to form a gradient that can be harnessed for activities like ATP synthesis (16). The retinal that ultimately enables the gradient is derived from carotenoids, colorful molecules that act as photoprotectants, cofactors, pigments, and membrane stabilizers throughout nature (17, 18). The initial building blocks of carotenoids are the five-carbon monomers isopentenyl diphosphate and dimethylallyl diphosphate. These units can be polymerized and modified to form lycopene, β-carotene, and various intermediates (19). Moreover, more complex keto-carotenoid chromophores like salinixanthin and echinenone boost rhodopsin activity by resonance energy transfer to retinal in rare xanthorhodopsin-like systems (20–22). Microorganisms with the machinery to produce functional, chromophore-loaded rhodopsins may have a competitive advantage in their native ecosystems (23, 24).

Here we provide experimental evidence that acI has photoheterotrophic potential. A significant challenge when these studies were initiated was the inability to purely culture acI (5, 25), although it appears this hurdle may soon be overcome (26). Consequently, we pursued a multidisciplinary approach that did not rely on large amounts of acI biomass, but instead focused on bioinformatic, transcriptomic, and biochemical data. Specifically, we characterize a biosynthetic pathway for retinal and its carotenoid precursors using heterologous expression. We then demonstrate that *acI-actR* is highly transcribed in the freshwater acI biome, Lake Mendota, and acI-ActR can function as a retinylidene holoprotein to produce a light-driven proton gradient. Most importantly, the nanomachine is active in cells that produce retinal using the native acI β-carotene oxygenase. Finally, we provide evidence for a complex carotenoid that may screen cells from oxidative damage and/or boost the light harvesting ability of actinorhodopsin.

## RESULTS

### All acI-ActR contain key amino acids for function

We first determined whether acI-ActR sequences were consistent with acI-holo-ActR formation. Indeed, all contain the features for proper rhodopsin structure and function (Fig. 1A). Seven predicted helices (α1-7) match those found in xantho-opsin, a close homolog of acI-ActR from *Salinibacter ruber* (20). A conserved lysine for Schiff base formation and acidic residues for proton-shuttling across the retinylidene gate are present (27), and a leucine is predicted to tune the absorbance of all acI-holo-ActRs to the green region (28). We also found novel features. A proline in the middle of α4 differentiates ActRs from xantho-opsins, and the residue may serve as a means for better phylogenetic classification (Fig. S1). Additionally, 3D structure predictions suggest cysteines on α1 and α7 in acI clades A and B form a disulfide staple (Fig. 1B). acI-ActRs also contain glycine near the top of α5, a required feature for binding the ketolated antenna carotenoids (21, 22). This glycine replaces a bulky residue, usually tryptophan, found in most rhodopsins, including the well-characterized bacteriorhodopsin.

**FIG 1.**
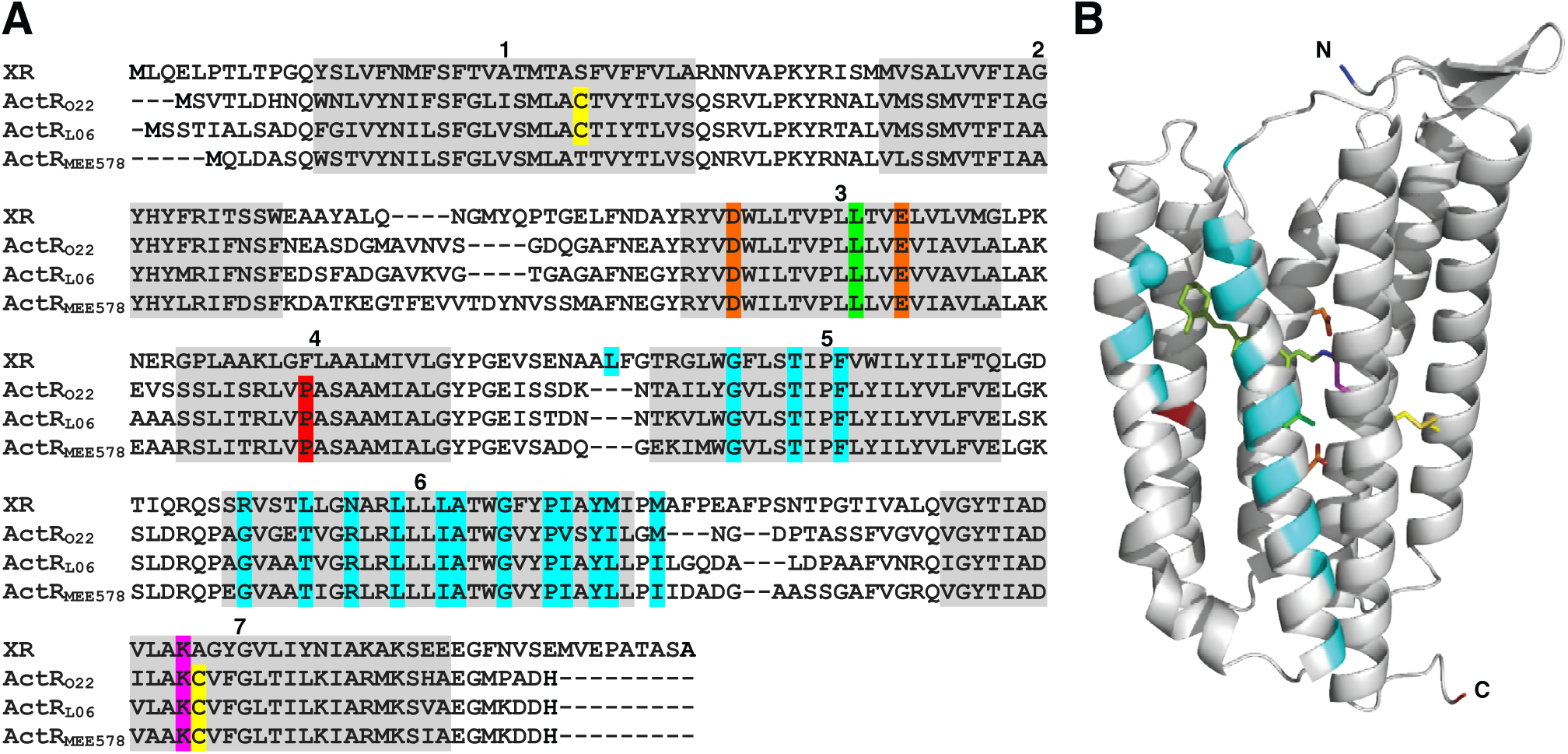
Features of acI-ActR from acI clades A, B, and C. (A) acI-ActR from clade A (ActR_O22_), clade B (ActR_L06_), and clade C (ActR_MEE578_) are aligned to a homolog, xantho-opsin (XR) from *Salinibacter ruber*. Proteins are subscripted with genome shorthand (Table S1). Features are highlighted: predicted helices (gray, numbered), cysteines (yellow), main proton shuttles (orange), main absorbance tuner (green), Schiff base lysine (magenta), possible antenna carotenoid residues (cyan), and proline indicative of ActR *vs* xantho-opsin (red). Antenna carotenoid residues are based on amino acids in proximity to salinixanthin in the crystal structure of xanthorhodopsin (PDB 3DDL). (B) 3-D structure prediction for ActR with key residue side chains displayed according to the color coding in panel (A). The α-carbon of glycine near the top of α5 that would allow antenna binding is shown as a sphere, the Schiff base retinal is modeled (lime green), the predicted disulfide bond of ActR from clades A and B is modeled, and the N- and C-termini are labeled.

### Seven gene products form an actinorhodopsin pathway

Retinal is the chromophore needed to form holo-ActR. We find that in addition to encoding ActR, acI single-cell amplified genomes (SAGs), metagenome-assembled genomes (MAGs), and complete genomes from dilution-to-extinction cultures (Table S1) appear to encode genes for enzymes that produce retinal and its carotenoid precursors (Fig. 2).

**FIG 2.**
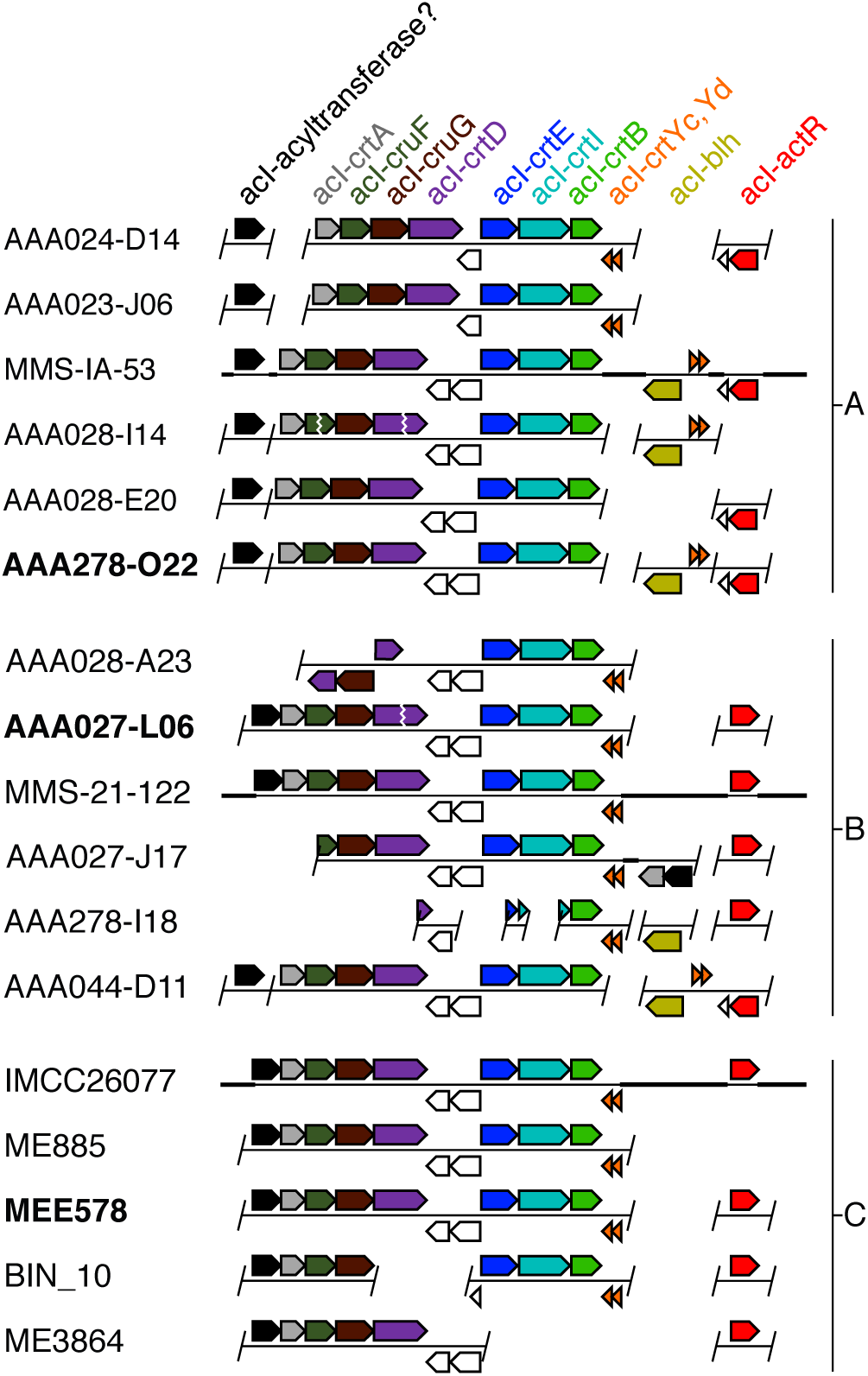
acI carotenoid-related genes in the genomic context. acI genomic contigs or genomes from clades A, B, and C are primarily labeled by shorthand notation (i.e. designation after “actinobacterium SCGC” or “int.metabat.”). Genes are arrows pointing in the direction of transcription. A slash indicates a contig boundary, which may or may not end immediately after the pictured gene, a vertical zig-zag indicates contig ends that have been manually paired, and a horizontal thick bar indicates a longer region of DNA not represented. Uncolored genes are neighboring genes that may or may not be functionally associated with the carotenoid-related genes. Bolded labels indicate a gene source for this study. Relevant gene numbers for this study from AAA278-O22 in IMG are 3540 (*acI-11*), 10590-10560 (*acI-crtA* to *acI-crtD*), 10530-10510 (*acI-crtE* to *acI-crtB*), and 7000-6980 (*acI-crtYd* to *acI-blh*), 1360 (*acI-actR*).

We assembled a plausible, complete pathway with protein assignments for forming lycopene, β-carotene, retinal, and subsequently, actinorhodopsin (Fig. 3A). Lycopene synthesis in acI requires three major steps: synthesis of geranylgeranyl-PP (Step 1), linkage of two geranylgeranyl-PP molecules to phytoene (Step 2), and tetra-desaturation of phytoene to lycopene (Step 3), predicted to be carried out by acI-CrtE, acI-CrtB, and acI-CrtI, respectively. Subsequent β-cyclization of lycopene would first produce γ-carotene (Step 4) then β-carotene (Step 5). A heterodimeric enzyme composed of acI-CrtYc and acI-CrtYd likely carries out these serial cyclizations, much like *Myxococcus xanthus* β-cyclase genes (36% and 33% identity, trihelical transmembrane topology, PxE(E/D) catalytic motif) (Fig. S2) (29-32). The symmetric cleavage of β-carotene by a β-carotene oxygenase, acI-Blh, would then form retinal (Step 6). acI-Blh is a putative dioxygenase based on 27% sequence identity, predicted helical topology, and four histidines for non-heme iron coordination that are shared with the characterized β-carotene dioxygenase from an uncultured marine bacterium (Fig. S3) (30, 31, 33). To form functional holo-ActR, retinal autocatalytically forms a Schiff’s base with the side chain of a conserved lysine in acI-ActR (Step 7).

**FIG 3.**
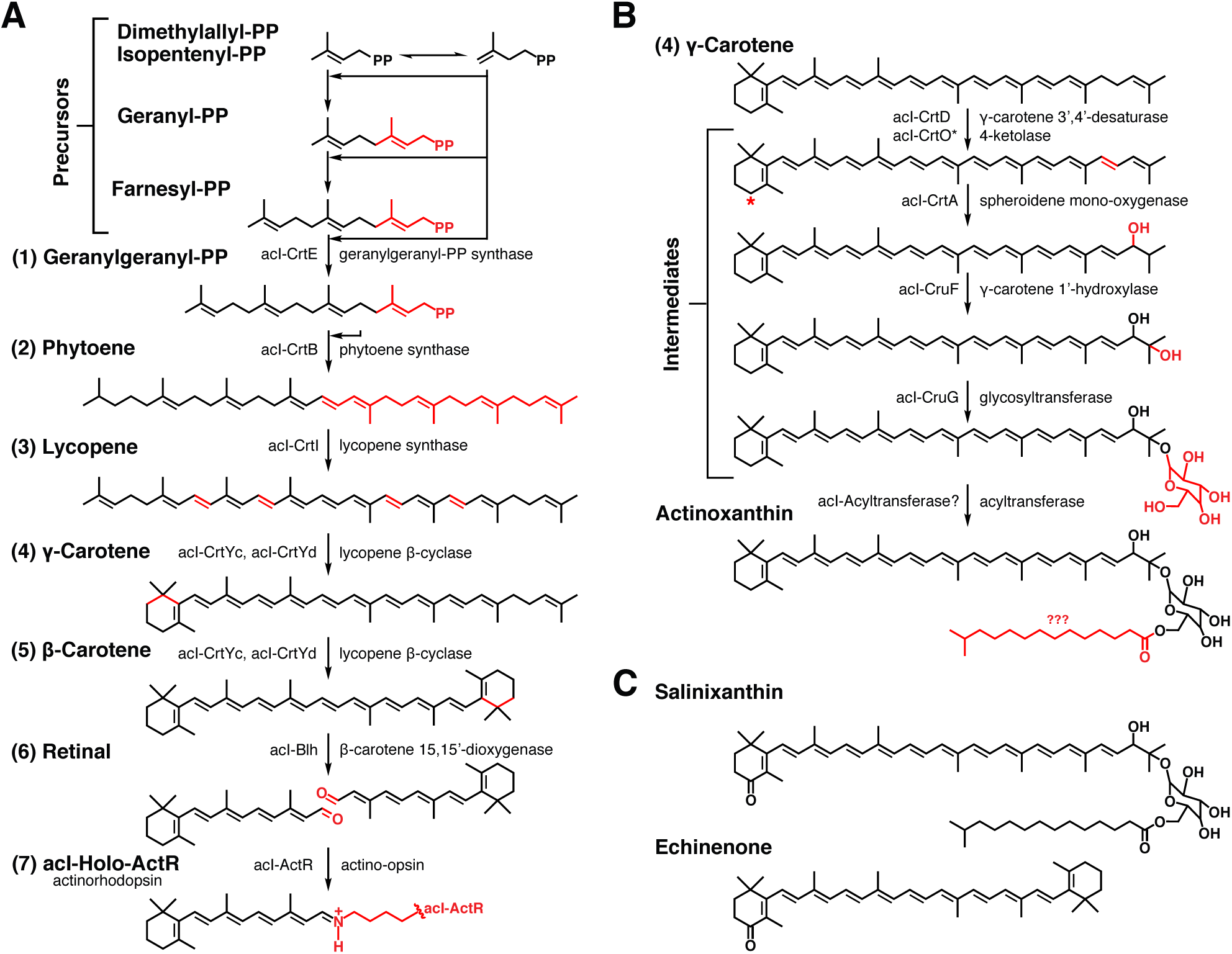
Predicted carotenoid-related pathways in acI. (A) The actinorhodopsin synthesis pathway requires isopentenyl precursors to be assembled into carotenoids (1-3), which are modified (4-5) and cleaved to produce retinal (6). A Schiff base forms between retinal and a lysine of acI-ActR to form acI-holo-ActR (7). (Chemical changes are shown in red, proteins from this study with their predicted functions are shown to the sides of progress arrows, and product names are shown to left of their chemical structures.) (B) A complex carotenoid synthesis pathway is predicted to require γ-carotene produced in the actinorhodopsin pathway with further desaturation, hydroxylation, glycosylation, and acylation of the carotenoid. Question marks indicate uncertainty in enzyme naming or chemical structure, and the asterisk represents the position of a carbonyl group if acI-CrtD is actually acI-CrtO, a ketolase. (C) Complex carotenoids from other organisms that function as carotenoid antennae on proton-pumping rhodopsins.

In the genomic context, genes encoding the enzymes for chromophore production are grouped into functional regions (Fig. 2). Lycopene production genes (*acI-crtE, acI-crtB, acI-crtI*) are found as a neighborhood in the order of steps 1, 3, 2 in all acI clades. Genes for steps 4-6 (*acI-crtYc, acI-crtYd, acI-blh*) are found at varying distances from the lycopene-synthesis cluster. In many cases, the 3’ ends of *acI-crtB* and *acI-crtYd* are adjacent but encoded in the opposite sense, thus forming a β-carotene-synthesis neighborhood. In other cases, the cyclase genes are instead found near *acI-blh*. In most cases, *acI-actR* is widely separated from other pathway genes. An interesting exception is AAA044-D11, where *acI-actR, acI-crtYc, acI-crtYd*, and *acI-blh* are contiguous (Fig. 2).

### A pathway for a complex carotenoid contains at least nine genes

We previously noted that genes for production of a predicted carotenoid glycoside-ester may be encoded adjacent to the lycopene gene neighborhood (3) (Fig. 2). In this study, we used bioinformatics and sequence homology to assign functions to gene products and assemble a plausible pathway (Figs. 3B).

The crucial carotenoid for initiating the pathway is γ-carotene (Step 4), and it is predicted to be produced in the retinal pathway. Modifications of γ-carotene are expected to include: desaturation of a carbon-carbon bond at the noncyclized end by acI-CrtD, introduction of two hydroxyl groups in separate steps by acI-CrtA and acI-CruF, and addition of glucose onto the terminal hydroxyl group by the glycosyl transferase, acI-CruG. Transfer of a fatty acid of unknown structure to the sugar by acI-acyltransferase would be the final step in the production of a novel carotenoid uniquely found in acI Actinobacteria. We propose to name this molecule actinoxanthin.

Genes encoding enzymes for actinoxanthin synthesis (*acI-crtA, acI-cruF, acI-cruG, acI-crtD*) are found upstream of the lycopene neighborhood as a contiguous group, which most often includes *acI-acyltransferase* (Fig. 3). Functional assignments for acI-CrtA, acI-CruF, acI-CruG, and acI-CrtD are based on sequence homology and topological predictions matching more well-characterized enzymes (Fig. S4, S5, S6, S7) (30, 31, 34–40). Because many carotenoid modification proteins are related in protein sequence (i.e. lycopene desaturases and carotene ketolases) (41), functional validation will be required for all of the carotenoid pathway enzymes. Specifically, the *acI-CrtD* gene product could encode a ketolase, *CrtO*, which would introduce a carbonyl onto the β-ionone ring rather than desaturating an additional bond (Fig. 3B). The resulting carotenoid would be structured like known rhodopsin antennae, salinixanthin and echinenone (Fig. 3C) (20–22). The acyltransferase identity and chemistry are also not well-defined, but the gene product’s involvement in actinoxanthin synthesis is supported by database annotation as a CoA methyl esterase.

### Metatranscriptomic analysis of pathway gene transcripts in environmental acI populations

For actinorhodopsin assembly in acI cells, *acI-actR* and retinal synthesis genes must be expressed. To measure gene expression in environmental acI, four metatranscriptome samples were collected across multiple time points from the surface of eutrophic Lake Mendota (Dane County, WI, USA) and sequenced. The resulting transcripts were mapped to available acI SAGs and MAGs (Table S1) to quantify relative gene expression levels in acI cells. Notably, *acI-actR* is the most highly expressed acI-A gene, the second most highly-expressed acI-B gene (11), as well as the most highly expressed gene from either pathway in each of the three acI clades (Table 1). Transcription is also significant for other genes in both the retinal and putative actinoxanthin pathways, but at levels 300 to 1000-fold lower than for *acI-actR* (Table 1). No transcripts were mapped for *acI-blh* from acI-C because the gene is neither present in any acI-C SAG or MAG genome nor the first complete acI-C genome obtained by dilution to extinction cultures (4).

**TABLE 1.** Log2 average reads per kilobase of transcript per million mapped read values for carotenoid-related gene transcripts

| Gene Name | *acl-A* | *acl-B* | *acl-C* |
| --- | --- | --- | --- |
| *acl-crtE* | 11.19 | 11.76 | 9.16 |
| *acl-crtB* | 9.33 | 9.64 | 9.23 |
| *acl-crtI* | 11.21 | 12.92 | 9.46 |
| *acl-crtYc* | 10.44 | 12.62 | 9.76 |
| *acl-crtYd* | ND<sup>a</sup> | 10.23 | 7.22 |
| *acl-blh* | 7.30 | 8.48 | NA<sup>b</sup> |
| *acl-actR* | 19.54 | 19.79 | 17.99 |
| *acl-crtD* | 9.39 | 11.98 | 8.29 |
| *acl-crtA* | 10.46 | 8.53 | 8.97 |
| *acl-cruF* | 9.80 | 10.28 | 9.09 |
| *acl-cruG* | 9.14 | 10.75 | 8.31 |
| *acl-11* | 9.23 | 9.04 | 7.93 |
| *recA* | 14.52 | 16.92 | 12.70 |

<sup>a</sup> Not Determined: Gene is present in clade member assemblies but no transcript mapped.
<sup>b</sup> Not Applicable: Gene is not present in any clade member assembly.

This mapping of Lake Mendota metatranscriptome samples provides direct evidence for actinoxanthin, lycopene, and lycopene cyclase operons (Fig. 4). Specifically, reads that mapped to the most populous genome, ME885, overlap the intergenic regions within each of the three operons (Figs. 3, 4). Synteny across acI clades, especially for the lycopene and actinoxanthin operons, similarly supports assignment of the three neighborhoods as genuine operons. Reading frame overlaps with *acI-crtA* in acI-C support *acI-acyltransferase* involvement in actinoxanthin synthesis.

**FIG 4.**
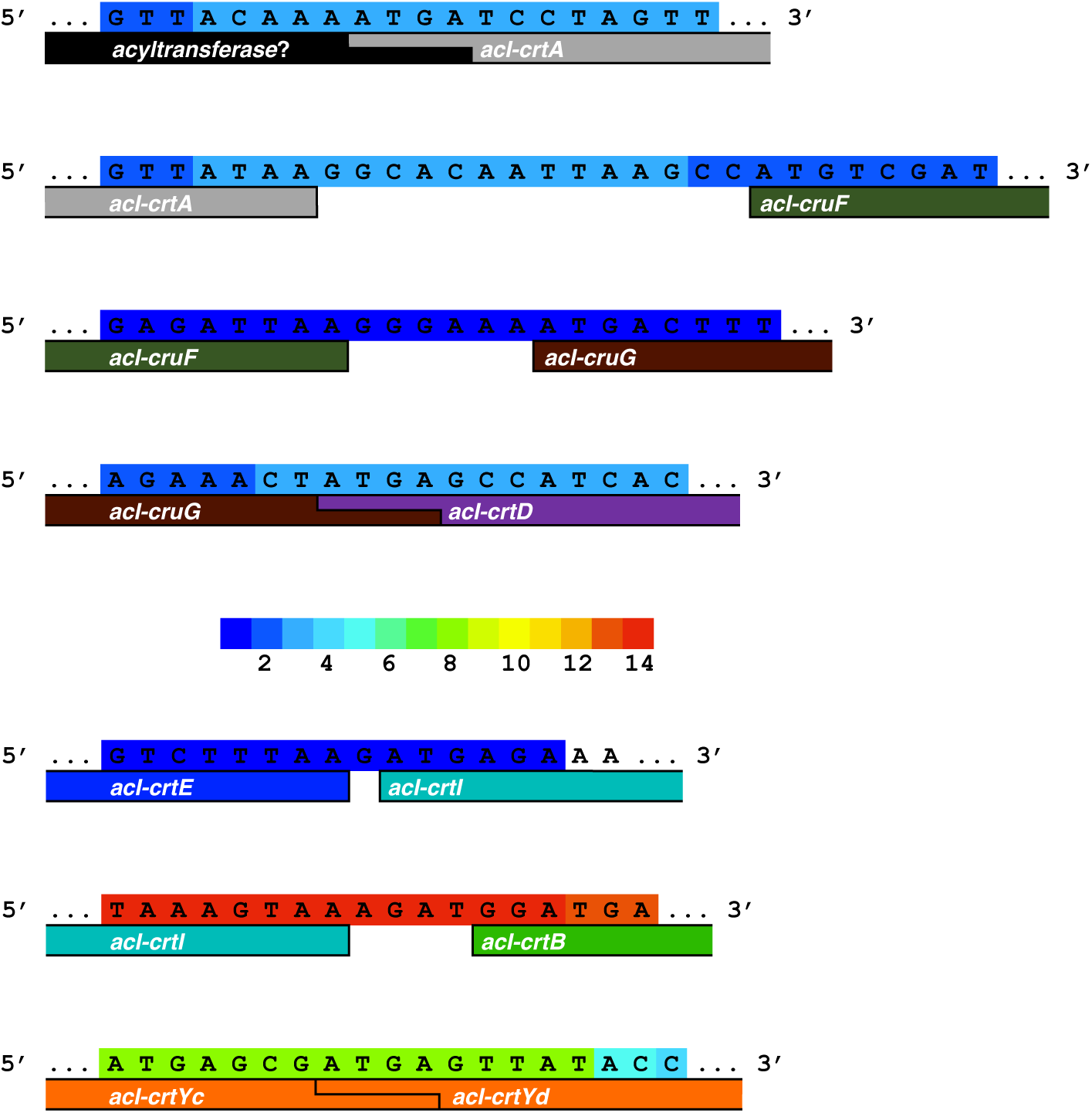
Intergenic transcripts that map to carotenoid-related genes in genome ME885. Operons are evident for actinoxanthin (*acyltransferase* through *acI-crtD*), lycopene (*acI-crtE* through *acI-crtB)*, and lycopene cyclase (*acI-crtYc* and *acI-crtYd*) biosynthesis. The color key in the center indicates of the number of times a base was covered. Transcripts and genes may continue beyond the edges of the DNA window.

### acI-CrtE, acI-CrtB, and acI-CrtI produce lycopene

We sought to demonstrate whether acI-CrtE, acI-CrtB, and acI-CrtI enzymes can form lycopene as predicted (Fig. 3A, Steps 1-3). Therefore, a fast, reproducible method for chromophore extraction from whole *E. coli* cells and identification by HPLC-mass spectrometry analysis was formulated. Relevant genes (*acI-crtE, acI-crtB*, and *acI-crtI*) from AAA278-O22 were cloned into pCDFDuet1 and expressed from the resulting acI-CrtEBI/-cassette (Table 2). Assembly AAA278-O22 was chosen for carotenoid production work because it contained all relevant genes and source DNA was available (3). The extracted compounds of these cells displayed absorbance maxima which exactly matched a lycopene standard at expected positions, 447, 472, and 504 nm (Fig. 5A) and displayed an intense red color (data not shown). HPLC-MS elution profiles depict lycopene geometric isomer peaks at m/z 536.438 (Fig. 5B). The retention time of the *all-trans* species (39.90 min) coincides for sample and standard as the peak with highest intensity. The lycopene assignment was further confirmed by MS/MS fragmentation of the parent ion.

**TABLE 2.**
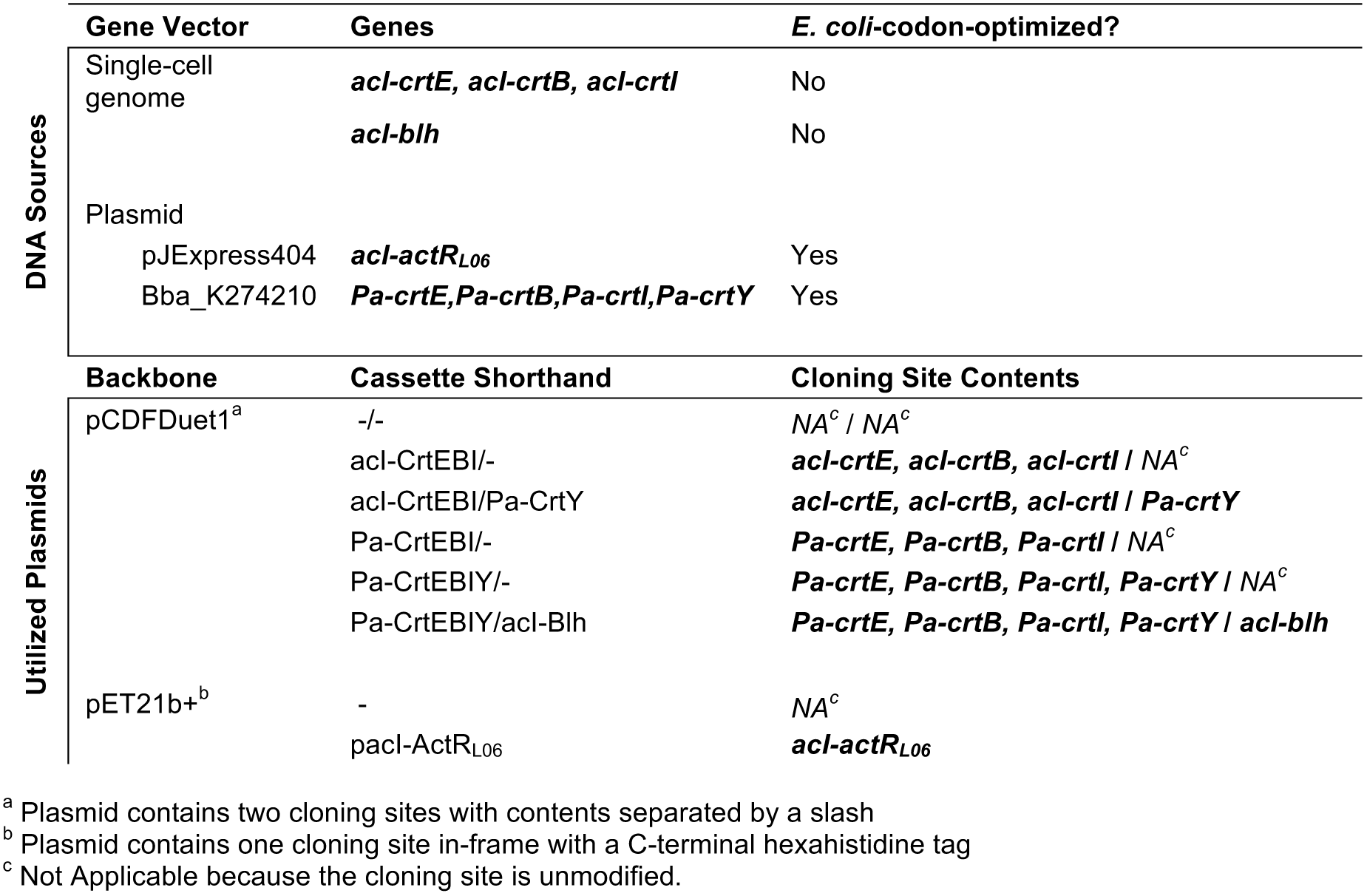
DNA sources and utilized plasmids for this study

**FIG 5.**
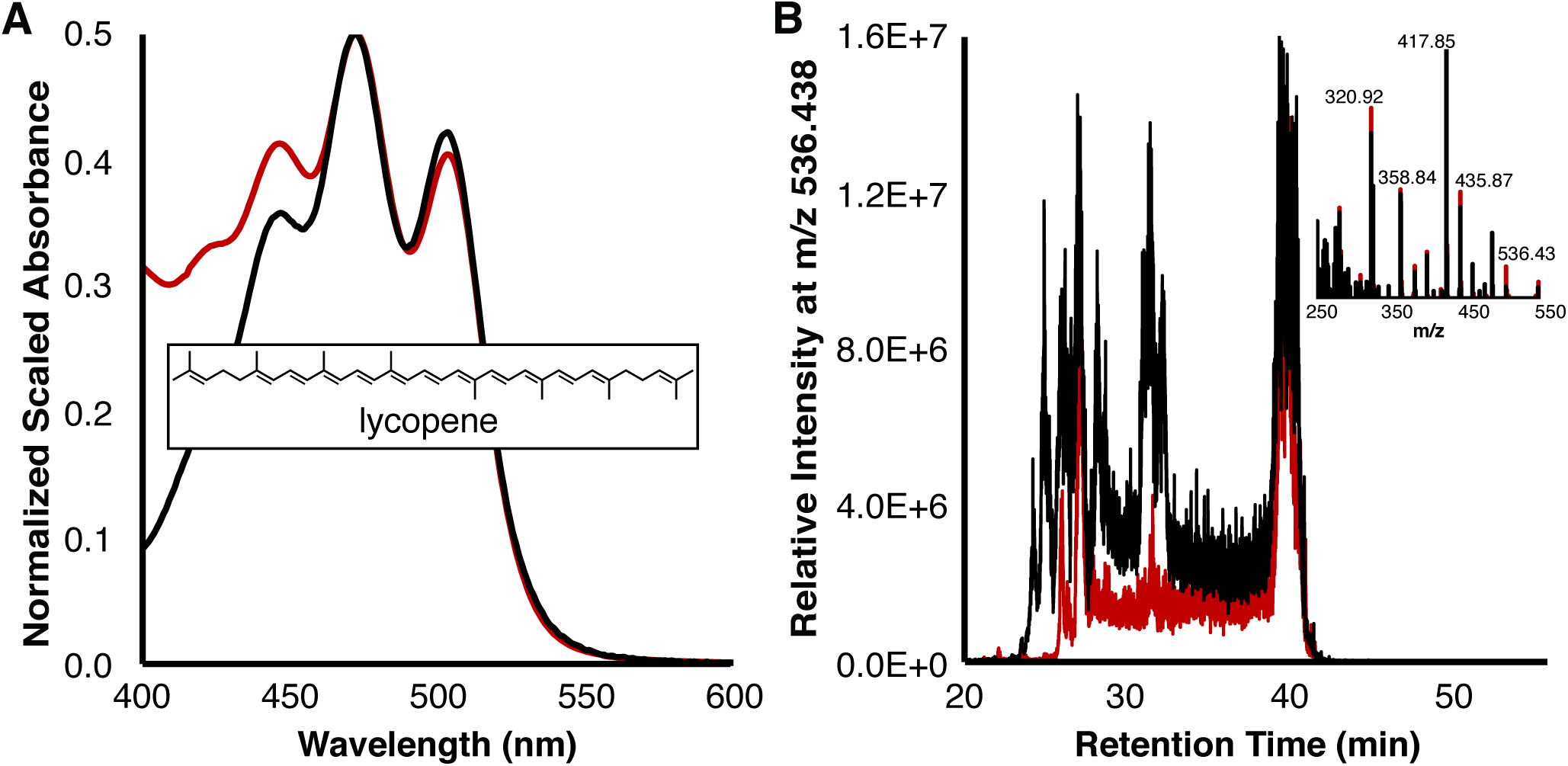
acI-CrtEBI catalyze lycopene formation. (A) Difference spectra for extracts from cells containing acI-CrtEBI/- *vs* -/- (black) and lycopene *vs*. acetone (red). The inset depicts lycopene. (B) HPLC-MS data for the same samples. The inset depicts MS/MS data from *all-trans* peak retention times.

To further characterize the lycopene product of acI-CrtEBI enzymes, we tested whether it serves as the substrate in a lycopene cyclization reaction. *Pantoea ananatis* genes *Pa-crtE, Pa-crtB, Pa-crtI* and/or *Pa-crtY* were expressed in *E. coli* as positive controls (Table 2, Fig. 6A). When the *Pa-crtY* cyclase gene and acI lycopene pathway genes were co-expressed from pCDFDuet1 acI-CrtEBI/Pa-CrtY, the cellular extract absorbance maxima were 407, 429 and 453 nm (Fig. 6B). These dramatically blue-shifted maxima, compared to β-carotene, combined with the defined cyclase activity of Pa-CrtY, identify the major extract product as β-zeacarotene, γ-carotene saturated between C7’ and C8’ (Fig. 6B). Thus, acI-CrtEBI produce a chromophore which serves as a substrate for Pa-CrtY lycopene cyclase. Presumably, β-carotene would be produced after proper desaturation and a second cyclization, which were inefficiently carried out in our non-native experimental set up.

**FIG 6.**
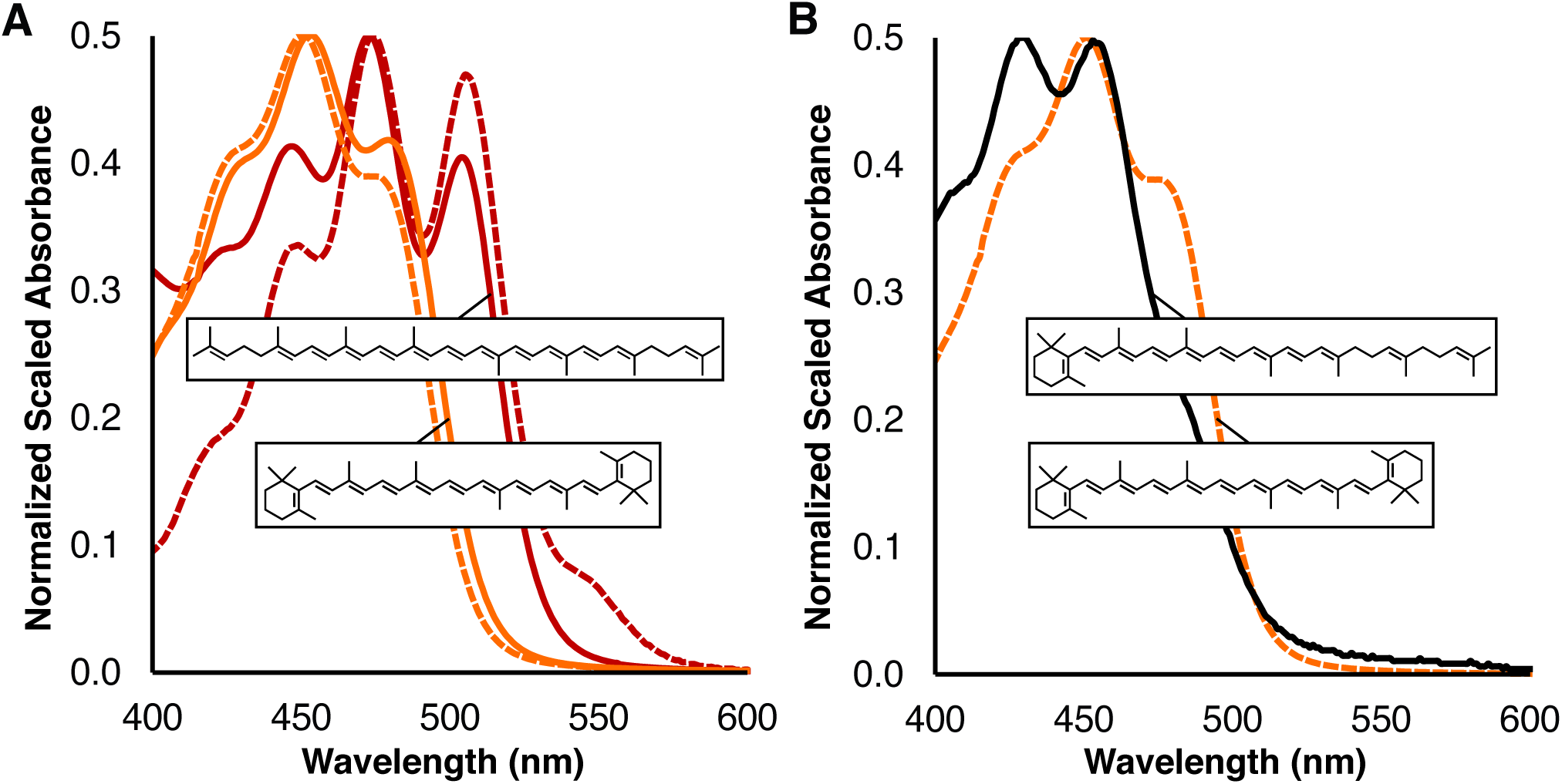
acI-CrtEBI-produced lycopene is cyclized by Pa-CrtY. (A) Difference spectra for Pa-CrtEBI/- derived lycopene *vs* -/- extract (red dash), standard lycopene *vs* acetone (red), Pa-CrtEBIY/- derived β-carotene *vs* -/- extract (orange dash), and standard β-carotene *vs* acetone (orange). The insets depict lycopene and β-carotene. (B) Difference spectra for synthesized β-carotene extract (orange dash) and acI-CrtEBI/Pa-CrtY *vs* -/- extracts (black). The insets depict β-zeacarotene and β-carotene.

### acI-Blh produces retinal from β-carotene

The final enzymatic step in retinal biosynthesis is the symmetric cleavage of β-carotene (Fig. 3A, Step 6). This enzyme is not encoded in any acI-C genomes (Fig 2). To show that acI can natively perform β-carotene cleavage, we tested the enzymatic activity of AAA278-O22 Blh. This *acI-blh* was expressed from pCDFDuet1 Pa-CrtEBIY/acI-Blh (Table 2) to ensure a large β-carotene substrate pool. Introduction of acI-Blh yielded yellow instead of intensely orange-colored cells observed when β-carotene is abundant (data not shown). HPLC-MS confirmed the presence of retinal at m/z 285.221 as well as a diagnostic retinol species at m/z 269.226 (Fig. 7A, 7B). As is the case for lycopene, geometric isomer peak patterning is observed. Maximal *all-trans-species* retention times match retinal and retinol standards at 23.76 min and 23.88 min, respectively. Additionally, MS/MS fragmentation of the appropriate m/z species confirmed retinoid identities (Figs. 7A, 7B). Retinol was roughly 7-times more abundant than retinal in the extract as judged by MS intensities. Accordingly, the UV/Vis absorbance profile appears more like retinol (Fig. 7C). Retinal produced by acI-Blh is likely serving as a potent electron acceptor for the *E. coli* alcohol dehydrogenase, *ybbO* (42, 43).

**FIG 7.**
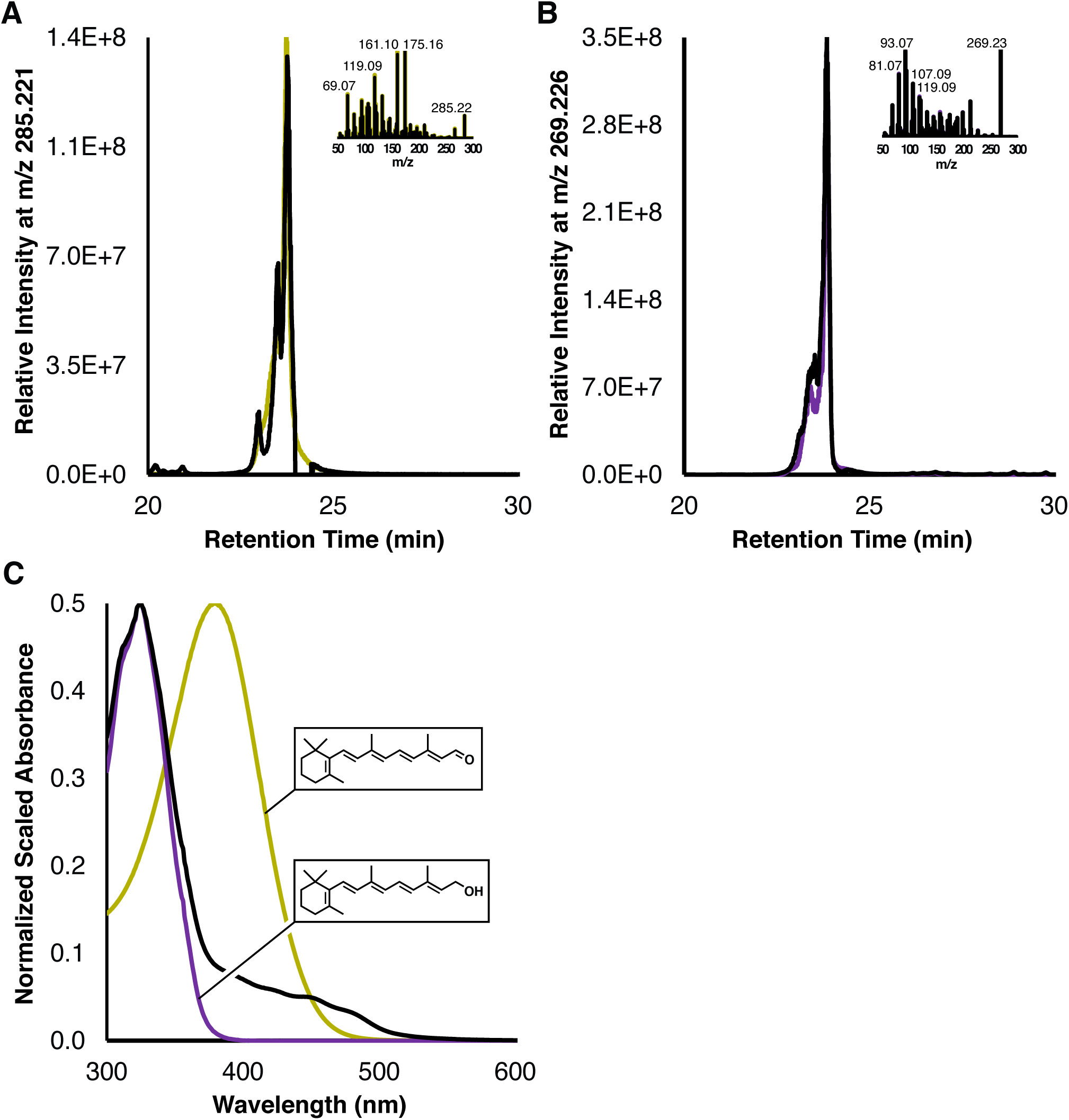
acI-Blh is a β-carotene oxygenase that catalyzes retinal formation. (A) HPLC-MS data for Pa-EBIY/acI-Blh extract (black) and a retinal standard (yellow). The inset depicts MS/MS data from *all-trans* peak retention times. (B) HPLC-MS and MS/MS (inset) data for the same experimental extract but with standard retinol spectra (purple). (C) Difference spectra for Pa-CrtEBIY/acI-Blh *vs* -/- (black) and retinal (yellow) or retinol (purple) standards *vs* ethanol. Insets depict retinal and retinol to highlight conjugation length differences.

### acI-ActR_L06_ with retinal forms an active, green light-dependent proton pump

The gold-standard test of a functional rhodopsin is light-dependent activity. Given that ActR proteins contain the residues for chromophore binding and proton movement, we tested an opsin from each clade (AAA278-O22, AAA027-L06, MEE578) for production and chromophore binding in *E. coli*. Each opsin was expressed from pET21b+ alongside the plasmid confirmed for retinal production. acI-ActR_L06_ was selected for further characterization due to its high expression and efficient retinal binding, as judged from samples captured by metal affinity from detergent-solubilized membranes. acI-holo-ActR_L06_ maximally absorbed in the green region at 541 nm (Fig. 8A). The covalent attachment of retinal was confirmed by incubation with hydroxylamine hydrochloride, which frees the retinal and leads to production of retinal oxime.

**FIG 8.**
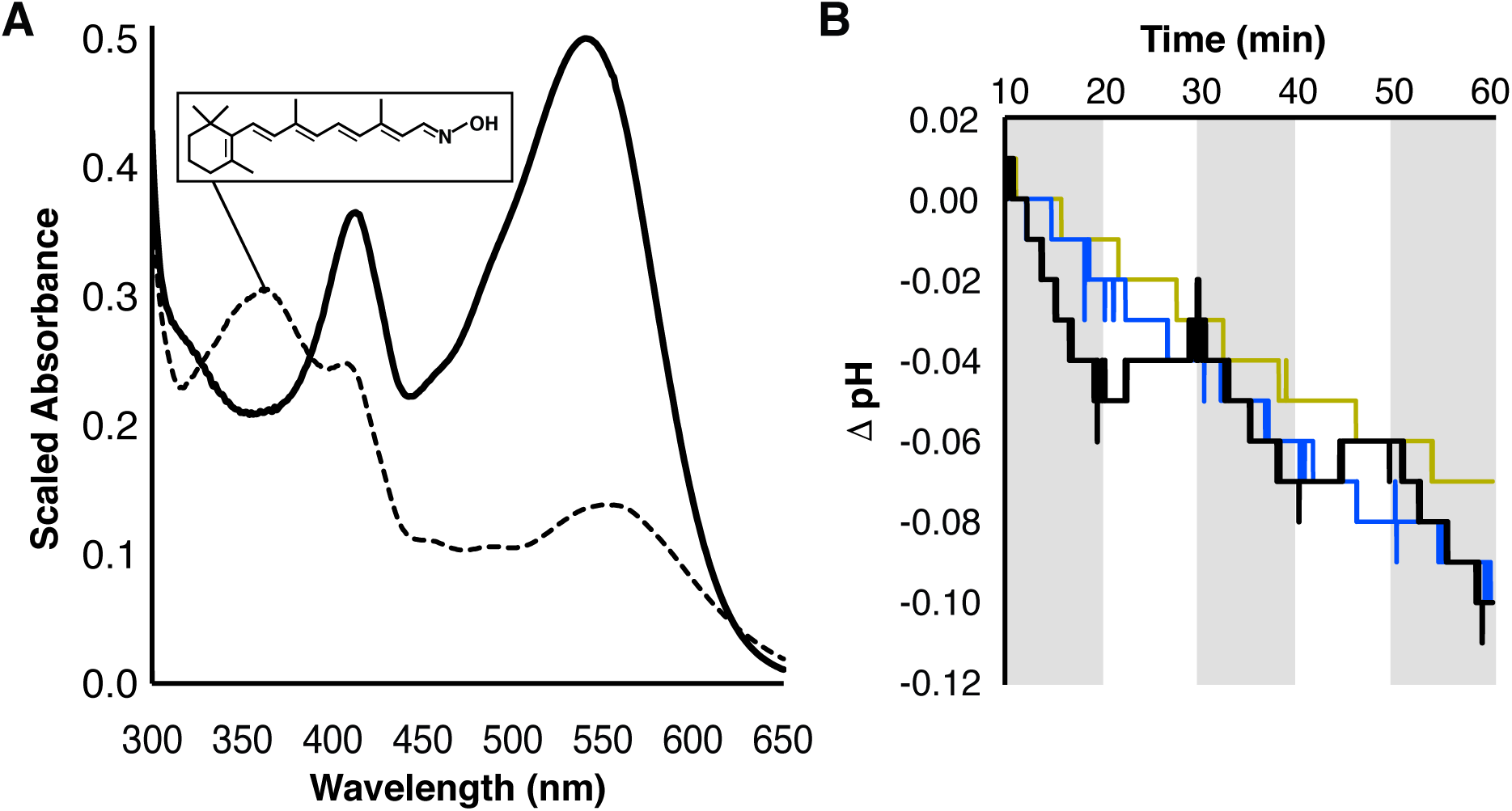
Holo-acI-ActR_L06_ is a light-driven proton pump. (A) Absorbance spectra for holo-acI-ActR_L06_ enriched from cells also expressing Pa-CrtEBIY/acI-Blh without (solid) and with (dashed) hydroxylamine hydrochloride. The retinal oxime that results from hydroxylamine incubation is shown. (B) Micro-electrode pH traces for cells expressing Pa-CrtEBIY/acI-Blh with (black, blue) or without (yellow) acI-ActR_L06_. CCCP is present in the blue trace. Shaded regions represent illumination by white light, and curves are from one of three independent experiments.

For conclusive demonstration that acI-holo-ActR_L06_ outwardly pumps protons in the presence of light when in a membrane environment, micro-electrode pH measurements were performed. During exposure to white light, the pH of a non-buffered assay solution sharply decreases compared to periods of ambient red light (Fig. 8B). To confirm protons as the source of the steep pH drop, carbonyl cyanide m-chlorophenyl hydrazone (CCCP) was added to discharge the proton gradient. CCCP trials did not display light-dependent proton-pumping but rather the constant downward drift of control cells. Thus, acI-holo-ActR is a retinal bound, green light-dependent proton-pumping rhodopsin.

## DISCUSSION

Prior to this study, there was no biochemical evidence to support the hypothesis that acI Actinobacteria use opsin-based phototrophy in freshwater. We have provided the first experimental exploration of the advantages that allow the acI lineage to be so abundant. We describe two favorable environmental adaptations, carotenoid biosynthesis and light-utilization. The former transforms simple isoprenoid precursors into carotenoids and retinal so that acI-ActR can function as a green-light-absorbing, outward proton-pumping acI-holo-ActR. The latter pathway is predicted to branch from a carotenoid intermediate in the acI-holo-ActR pathway to synthesize a complex carotenoid, actinoxanthin, with a yet unconfirmed structure and function.

The pathway for retinal and rhodopsin synthesis has been experimentally confirmed using acI Actinobacterial proteins. The machinery to start retinal production consists of three enzymes, acI-CrtE, acI-CrtB, and acI-CrtI (Steps 1-3). The enzymes produce lycopene (Fig. 3A, 5) and cluster into the *acI-crtEIB* operon (Figs. 2, 4). The reactions for γ-carotene and β-carotene (Steps 4, Step 5) that follow lycopene synthesis are likely carried out by a heterodimeric membrane protein expressed from the *acI-crtYc* and *acI-crtYd* operon (Figs. 2, 3, 4). Homologs of these gene products were found in *Myxococcus xanthus* and confirmed to synthesize a mix of γ-carotene and β-carotene (29). Therefore, we propose that acI can also synthesize these chromophores. Although the ratio of products in acI is unknown, it may be intrinsically linked to intracellular concentrations of retinal and actinoxanthin. Even lycopene formation may be a regulatory point for carotenoid synthesis in acI because coexpression of lycopene and lycopene cyclase does not guarantee dicyclized carotenoid (Fig. 6). Unsaturated but cyclized intermediates like β-zeacarotene may result from differing levels of cyclase compared to acI-CrtE, acI-CrtB, and acI-CrtI. Abundant cyclase may disrupt a substrate channeling assembly for lycopene as seen in acI-CrtEBI, Pa-CrtY-producing cells. Along these lines, an efficient co-localized assembly would help explain the discrepancy between extremely high levels of *actR* versus other pathway transcript levels in Lake Mendota (Table 1). We note that a system for stable β-zeacarotene production could prove a fruitful biotechnology tool, because monocyclized carotenoids are not readily available for purchase.

A key finding of our work is the presence, expression, and robust activity of a retinal-producing oxygenase from the acI gene, *acI-blh*. The enzyme symmetrically cleaves β-carotene to two retinal molecules (Step 6) (Figs. 3A, 7) and might additionally use alternative substrates such as a β-zeacarotene- or γ-carotene-like carotenoid to yield a single retinal. Notably, the *acI-blh* gene is not a feature of acI-C; in five acI-C genomes ranging in completeness from 25-100%, *acI-blh* has never been recovered (Table S1) (4). This absence points towards clade-level differentiation with respect to retinal production and subsequent acI-holo-ActR assembly. Specifically, we propose that tribes or populations of acI-A and acI-B with the ability to synthesize retinal display population persistence because of photoheterotrophy. This is in contrast to acI-C that follow a seasonal “bloom and bust” lifestyle possibly because of a reliance on exogenous chromophores (e.g. from cyanobacteria) (44), for example through uptake of retinal by *acI-blh-*lacking cells from lysed cyanobacteria. Many species of cyanobacteria encode functional β-carotene oxygenases (33, 45, 46), and retinoid concentrations in eutrophic lakes during cyanobacterial blooms are measurable (47). This proposal is consistent with actinorhodopsin studies in the culturable freshwater organisms *Rhodoluna lacicola* and *Candidatus* Rhodoluna planktonica (12, 13). While both encode *actR*, only the latter contains *blh* and thus exhibits self-sufficient rhodopsin activity in the laboratory. *R. lacicola* is dependent on retinal supplementation. As such, it may be analogous to acI-blh-lacking organisms. Indeed, recently reported complete genomes (5) indicate that even some acI-A and -B would require an alternative retinal source. This heterogeneity in gene content has major implications for niche-filling by acI-A and acI-B members and may dictate community-related growth patterns.

Regardless of where acI source retinal, all acI-ActR proteins contain the necessary features for proper activity as retinal holoproteins with green maximal absorbance, and we have demonstrated that an example acI-ActR functions as a retinylidene protein (Figs. 1, 8). Green absorption correlates with light penetration depth for many freshwater bodies where acI thrive, and strengthens the case that acI-holo-ActR is a light-activated protein in the environment. Native production of holo-ActR would have a major impact on energy availability for the acI cells. In addition to showing its proton-pumping activity, we discovered interesting qualities of acI-ActR (Figs. 1, S1). A proline in helix four was determined to be a phylogenetically differentiating residue between actino-opsins and other xantho-opsins, like the one in *Salinibacter ruber*. Structurally, prolines kink helices and the residue may have broad effects on photocycling times and/or intermediate photocycle structures. Similarly, a disulfide stapling of α1-α7 in acI-A and acI-B could also impact activity and/or protein stability. Additionally, it is interesting that acI-C organisms lack both the cysteine pair and *acI-blh*.

In addition to retinal and ActR synthesis, members of all acI clades contain an operon for other carotenoid biosynthetic machinery (*acI-acyltransferase, acI-crtA, acI-cruF, acI-cruG, acI-crtD*) (Figs. 2, 4). The gene products might produce actinoxanthin, a complex carotenoid with glycosyl and acyl modifications (Fig. 3B). Although the structure and presence of the carotenoid remain to be experimentally validated, the glycine void predicted to be near the top of α5 hints at possible secondary chromophore photoactivity. This variance from a bulky residue is necessary although not sufficient for binding of antenna keto-carotenoids (48, 49). This would be the third example of a special class of rhodopsins that we call antenna rhodopsins (20–22), but others are predicted (49). We are left with the question: can acI synthesize a complex carotenoid using the genes found upstream from the lycopene cluster, and does it interact with acI-ActR as an antenna?

Two biological scenarios for carotenoid-related light utilization in acI include: acI-holo-ActR is an antenna rhodopsin that augments proton-pumping via carotenoid energy transfer, or a complex carotenoid protects the cell from oxidative species and excess light while the proton-pump functions (Fig. 9) (50). Both situations favor production of a proton gradient to power numerous nutrient uptake transporters particularly when nutrients are limited (11). Such adaptations go far toward explaining how acI Actinobacteria can reach nearly 50% of the microbial cell number in freshwater lake epilimnia, and why they exist in virtually all freshwater ecosystems (2, 51). Further defining the physiology of a dominant ecological force within the lake biome will fill a gap in our conceptual models for freshwater carbon and energy cycling and improve ecosystem behavior forecasting.

**FIG 9.**
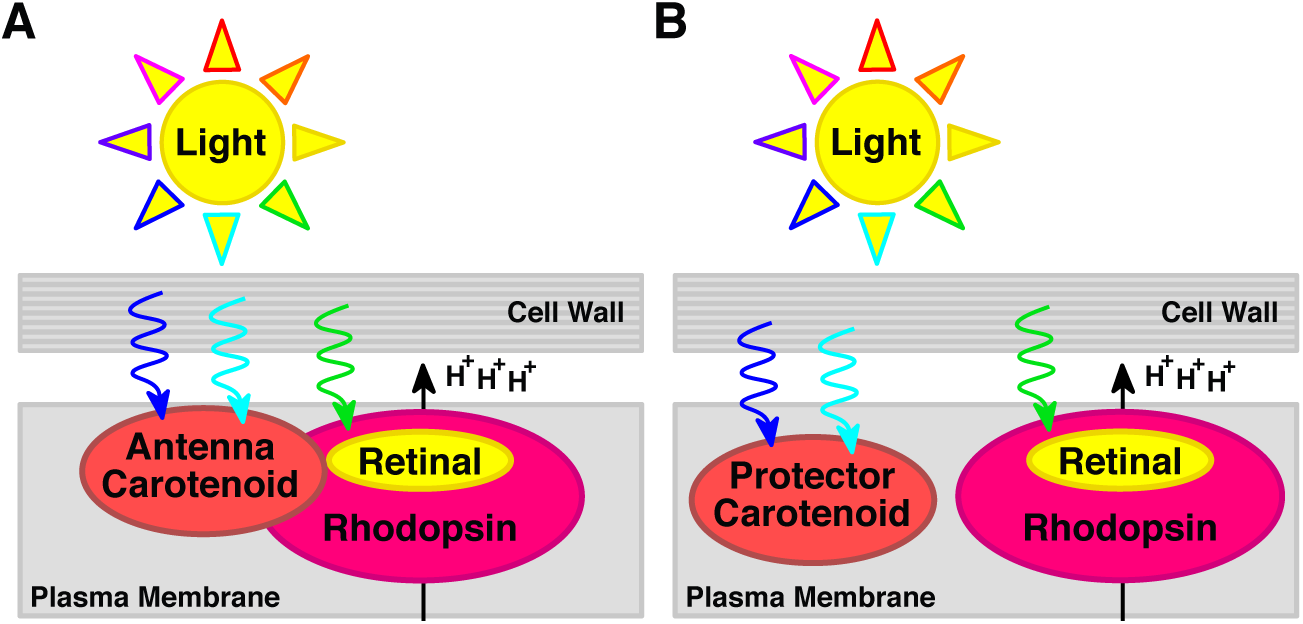
Biological model for acI actinobacterial carotenoid usage. (A) acI cells may utilize a complex carotenoid as an antenna that transfers light energy into the green-light-absorbing proton-pumping rhodopsin. (B) acI cells may utilize a complex carotenoid as protection from excess light and oxidative species to protect rhodopsin.

## MATERIALS AND METHODS

### acI Gene Identification and Pathway Assembly

Annotations for multiple acI SAGs (3) and MAGs were analyzed for genes relating to carotenoids using the Joint Genome lnstitute’s lntegrated Microbial Genome’s Viewer (52). Translated candidate protein sequences were used to identify homologs in other acI genomes, and those with consistent gene neighborhoods and known carotenoid-related functions were prioritized. The gene cassette with functional assignments was grouped into two pathways for carotenoid biosynthesis and use in acI Actinobacteria.

### acI Biomass Collection and Transcriptomics

Environmental sampling, metatranscriptome sequencing, and gene-expression calculations were all performed as previously described (11). Briefly, four samples were collected from within the top 12m of the Lake Mendota (Dane County, WI) water column, filtered through cheesecloth and onto 0.2 μm mixed cellulose filters (Whatman). Filters were immediately vitrified in liquid nitrogen and stored at −80°C. Samples were subject to RNA extraction, ribosomal RNA (rRNA) depletion, and sequenced on an Illumina HiSeq2500. All protocols and scripts for sample collection, RNA extraction, rRNA depletion, and sequencing are on GitHub (https:github.com/McMahonLab/OMD-TOIL).

Because this work used metatranscriptomes obtained as part of a previous study (11), raw paired-end reads were trimmed, merged, subject to *in-silico* rRNA removal, pooled, and mapped to a single reference FASTA file containing 36 high-quality acI genomes from a larger freshwater genome collection (11). Reads were competitively mapped to genes using BBMap (https:sourceforge.net/projects/bbmap/) and counted using HTSeq-Count (53). Raw count data were then used to compute gene expression on a Reads Per Kilobase of transcript, per Million mapped reads (RPKM) basis (54). All scripts are found on GitHub (https:github.com/joshamilton/Hamilton_acI_2017).

### Plasmid construction

All cloning was performed using Phusion HF GC Master Mix (ThermoFisher) and the listed primers (Table S2). A collection of stable DNA sources for amplification and subcloning was first created. Plasmid containing *E*. coli-codon-optimized *crtE, crtB, crtI, crtY* from *Pantoea ananatis* (Pa) was obtained from the International Genetically Engineered Machine (iGEM) organization catalog (http:parts.igem.org/Main_Page) (Table 2). The genes were PCR-amplified from the iGEM plasmid as a block. acI biosynthetic genes were PCR-amplified from AAA278-O22 genomic DNA as gene clusters. The L06 opsin sequence was obtained as an *E. coli*-optimized gene from DNA2.0.

All genes were individually PCR-amplified, if needed, and added to a cloning pipeline. Primer design included up to 40 base pairs of cloning site flanking regions from pCDFDuet-1 (Novagen). Genes were placed into the first site between NcoI and AflII restriction sites and between NdeI and AvrII sites of the second site. Appropriately restriction-enzyme-digested and gel purified plasmid backbone was transformed with up to a 10-fold molar excess of PCR product into *E. cloni* 10G cells (Lucigen) for ligation free plasmid recombination. After selection on 100μg/mL spectinomycin sulfate LB-Miller agar plates, insert presence was validated by colony PCR using GoTaq Green Master Mix (ThermoFisher). Restriction enzyme digestion in lab and Sanger sequencing at the UW Biotechnology Center further confirmed plasmid correctness. pET21b+ (Novagen) was used in the same pipeline ensuring genes were in-frame with a C-terminal hexahistidine tag with selection by 100μg/mL carbenicillin, sodium salt.

### Chromophore Production and Extraction

For chromophore production, multiple colonies of freshly transformed BL21 (DE3) Tuner *E. coli* (EMD Millipore) were picked into half-full flasks of LB-Miller broth plus 100μg/mL spectinomycin sulfate and shaken in 37°C darkness at 250 rpm for 24 hours. Five hundred or 1500 ODmL (ODmL = OD_600nm_* dilution * volume in milliliters) cells were centrifuged at 3300xg, washed in 100mM Tris 6.8, centrifuged again, and vitrified in liquid nitrogen. For chromophore extraction, cells stored at −80°C were thawed at 23°C for up to 30min. Cells were suspended at 500ODmL cells/3 mL acetone (Sigma, HPLC-grade), intensely vortexed for 1 min, and iced for 5 min. The chromophore containing supernatant after a 10 min 8000 x g clarification was added to 1mL/500ODmL of 23°C NaCl-saturated water (Fisher, ACS-grade) and 1mL/500ODmL 23°C dichloromethane (ACROS ACS-grade) and vortexed for 1 min. Further centrifugation resulted in a colorless aqueous bottom layer and a top colorful organic layer. The organic layer was evaporated under nitrogen, resulting solids were suspended in appropriate solvent, and filtered through a compatible 4mm 0.22μm filter.

### UV-Vis and LC-MS/MS analysis of chromophores

Absorbance spectra were acquired on a Beckman Coulter DU640B spectrophotometer. A Dionex Ultimate 3000 UHPLC coupled by electrospray ionization (ESI; positive mode) to a hybrid quadrupole - high-resolution mass spectrometer (Q Exactive orbitrap, Thermo Scientific) was used for detection of target compounds based on their accurate masses, mass spectra, and retention times (all matched to purified standards). Liquid chromatography (LC) was based on a published protocol (55). Separation was achieved using a YMC C30 reversed-phase carotenoid column (150mm × 2.5mm, 3μm particles) (SOURCE) at a flow rate of 0.2mL/minute. Solvent A consisted of equal parts methanol and water with 0.5% v/v acetic acid; Solvent B was equal parts methanol and methyl-tert-butyl-ether with 0.5% v/v acetic acid (55). Total run time was 58 min with the following two gradients: 5 min 30% B, 20 min ramp to 100% B, 100% B (retinoids) or 5 min 30% B, 25 min ramp to 100% B, 100% B (carotenoids). MS scans consisted of full positive mode scanning for m/z 200-600 from 5 min onwards. In addition, MS/MS scans were obtained by isolating fractions with m/z of 537.44548 (lycopene), 285.22129 (retinal), and 269.22639 (retinol). MS/MS fragmentations were performed at 30 normalized-collision energy (NCE) with an isolation window of 1.4 m/z and a post-fragmentation window of 50 to approximately 25 m/z above the isolation mass. For all scans, mass resolution was set at 35000 m/z ppm, AGC target was 1 × 10^6^ ions, and injection time was 40ms. Settings for the ion source were: auxiliary gas flow rate – 50, sheath gas flow rate – 10, sweep gas flow rate – 2, spray voltage – 3.5 kv, capillary temperature – 350°C, heater temperature – 250°C, S-lens RF level – 55.0. Nitrogen was used as nebulizing gas by the ion trap source. Standards for lycopene (Toronto Research Chemicals Inc., L487500, Lot 7ANR-20-1) β-carotene (Sigma Aldrich, PHR1239-3×100mg, Lot LRAA6761), retinal (Sigma Aldrich, R2500-25MG, Lot SLBN4199V), and retinol (Sigma Aldrich, R7632-25MG, Lot BCBP8066V) were analyzed along with experimental samples. Isomer peaks within the mass window result most often from environment-induced changes during sample preparation. Control cells did not display any significant sustained signal within the time-mass window. Data analysis was performed using Thermo Xcalibur and visualized on MAVEN (56, 57) and Thermo Xcalibur (Thermo Scientific) software.

### Proposed gene product and actino-opsin analysis

Clustal Omega was used to align proposed biosynthetic enzymes and opsin sequences for identification of protein characteristics (58–60). Opsin sequences were submitted to the I-TASSER server for structure prediction using xantho-opsin (PDB 3DDL:A) as the template (61–63). Basic Local Alignment Search Tool (BLAST) and TM/HMM 2.0 were used for identity percentage calculation and transmembrane helix estimates, respectively (30, 31). PyMOL was used to visualize the resulting protein structures (64).

The actino-opsin phylogeny was reconstructed using opsin protein sequences from bacterial isolate references, SAGS and MAGS. Opsin protein sequences were aligned with PROMALS3D multiple sequence and structure alignment tool (65). The alignment was trimmed to exclude columns that contained gaps for more than 30% of the included sequences. Poorly aligned positions and divergent regions were further eliminated by using Gblocks (66). A maximum likelihood phylogenetic tree was constructed using PhyML 3.0 (67), with the LG substitution model and the gamma distribution parameter estimated by PhyML and a bootstrap value of 100 replicates. The phylogenetic tree was visualized with Dendroscope (v3.2.10) with midpoint root (68). The ActR and other XR sequence subtree was extracted and displayed.

### Actinorhodopsin enrichment and analysis

BL21 (DE3) Tuner *E. coli* cells co-transformed with plasmid expressing Pa-CrtEBIY/acI-Blh and acI-ActR_L06_ were grown as during carotenoid production except that carbenicillin, sodium salt was also added to 100μg/mL. All further steps were done under red light or darkness at 4°C and/or on ice using 4°C buffers. 2000 ODmL cells were harvested and suspended in 5g/mL Lysis Buffer (50mM Tris-HCl 8.0, 300mM NaCl, 1 50mL EDTA-free protease inhibitor tablet (Roche), 1mg/mL lysozyme, 20μg/mL DNase 1, 5mM MgCl_2_, 130μM CaCl_2_, 4mM PMSF). Cells were lysed at 16000 psi by five passes through a French Pressure cell. Sequential centrifugation at 10,000xg for 15 min and 100,000xg for 45 min cleared debris and pelleted membranes. Membranes were loosened with Lysis Buffer and transferred to a Potter-Evelhjem homogenizer. After homogenization, membranes were diluted with 4°C Lysis Buffer to 25mL and recentrifuged at 100,000xg for 45 min. Suspension and homogenization were repeated with Solubilization Buffer (50mM Tris-HCl 8.0, 300mM NaCl, 2% m/v dodecyl-maltoside, 10mM imidazole). Membranes were diluted to 15mL with Solubilization Buffer and rocked 18hr in darkness. Centrifugation at 20000xg for 20min clarified the material before chromatography. The soluble fraction was loaded at 0.5mL/min onto an equilibrated 1mL Ni-NTA column. The processing profile in column volumes (CV) was: 3 Solubilization Buffer, 12 Wash Buffer (50mM Tris-HCl 8.0, 300mM NaCl, 0.05% m/v DDM, 30mM imidazole), 5 Elution Buffer (50mM Tris-HCl 8.0, 300mM NaCl, 0.05% m/v DDM, 500mM imidazole). The first elution fraction was dialyzed against 1L 4°C Final Buffer (10mM HEPES 7.5, 100mM NaCl, 0.05% m/v) for 2hr and then analyzed by absorbance spectroscopy. Light and 50 μL 23°C saturated hydroxylamine HCl was used to selectively remove retinal from 140 μL of sample to confirm the Schiff base. The resulting unbound retinal oxime absorbs at wavelength maxima between that of retinal and retinol where 247 and 257 nm peaks indicate 11-cis and all-trans oxime species, respectively.

### Micro-electrode pH trace acquisition

The assay was similar to a published protocol and uses *E. coli* for expression of holo-ActR (12). All steps were carried out under red light generated by putting red cellophane over a fluorescent bulb. 500 ODmL *E. coli* cells were harvested by centrifugation at 3300xg for 15 min at 4°C. Cells were suspended in 45 mL 23°C Assay Solution (10mM NaCl, 10mM MgSO_4_-7H_2_O, 100uM CaCl_2_-2H_2_O) and centrifuged at 3300 x g for 10 min. The latter wash step was repeated. Final suspension was in 20 mL 23°C Assay Solution to yield an OD=25. Cells were incubated for 60 min at 23°C to stabilize in darkness before the onset of the assay. The assay was set up with the sample in a glass test tube surrounded by a 300 mL 23°C water bath in a 400 mL beaker. A Mettler-Toledo InLab Micro probe (model number 51343160) was clamped in place above the sample tube and connected to a Sper Scientific Datalogger (model number 850060). The entire setup was surrounded by foil lined cardboard on four sides. Two FE15T8 bulbs (15W, 700 lumens each) in an 11×45 cm housing with 90° reflectors were placed 15 cm from the center of the sample tube such that light illuminated the entire length of the closest tube side. pH was recorded every second after 3.5 min of equilibration time for 60 min as follows: three cycles of 10 min with fluorescent lights off and 10 min fluorescent lights on. Carbonyl cyanide m-chlorophenyl hydrazone (CCCP) (Alfa Aesar, L06932 Lot 10181844) was added to a final concentration of 20 μM during dark stabilization after 45 min.

## ACKNOWLEDGEMENTS

This work was supported by the National Institute of Food and Agriculture, U.S. Department of Agriculture (2016-67012-24709 to JJH and WIS01789 to KDM), the United States National Science Foundation (NTL-LTER DEB-1440297 and DEB-1344254 to KDM, MCB-1518160 to KTF) and the National Oceanic and Atmospheric Administration (NA10OAR4170070 to KDM and KTF via the Wisconsin Sea Grant College Program Project #HCE-25). We thank former McMahon laboratory member, Dr. Sarahi Garcia, for the initial AAA0278-O22 DNA, former and current McMahon laboratory members for help obtaining transcript data, Diego Yanez for assisting with RNA methodology, and Dr. James Steele for supplying plasmid iGEM DNA.

